# Integrative Gene Expression and Metabolic Analysis tool IgemRNA

**DOI:** 10.1101/2021.08.02.454732

**Authors:** Kristina Grausa, Ivars Mozga, Karlis Pleiko, Agris Pentjuss

## Abstract

Genome scale metabolic modelling is widely used technique to research metabolism impacts on organism’s properties. Additional omics data integration enables a more precise genotype-phenotype analysis for biotechnology, medicine and life sciences. Transcriptome data amounts rapidly increase each year. Many transcriptome analysis tools with integrated genome scale metabolic modelling are proposed. But these tools have own restrictions, compatibility issues and the necessity of previous experience and advanced user skills. We have analysed and classified published tools, summarized possible transcriptome pre-processing, and analysis methods and implemented them in the new transcriptome analysis tool IgemRNA. Tool novelty is the possibility of transcriptomics data pre-processing approach, analysis of transcriptome with or without genome scale metabolic models and different thresholding and gene mapping approach availability. In comparison with usual Gene set enrichment analysis methods, IgemRNA options provide additional transcriptome data validation, where minimal metabolic network connectivity and flux requirements are met. IgemRNA allows to process transcriptome datasets, compare data between different phenotypes, execute multiple analysis and data filtering functions. All this is done via graphical user interface. IgemRNA is compatible with Cobra Toolbox 3.0 and uses some of its functions for genome scale metabolic model optimization tasks. IgemRNA is open access software available at https://github.com/BigDataInSilicoBiologyGroup/IgemRNA.

## INTRODUCTION

In the last decade the amount of different sequences has increased rapidly and counts more than 219 * 10^6^ (https://www.ncbi.nlm.nih.gov/genbank/statistics/ on 06.05.2021). Escalating genome availability, recent discoveries and technological advancements in sequencing and mass spectrometry have caused an increase in multi - omics data set generation. From a pure mathematical and statistical point of view multi-omics data set analysis is still very challenging and lacks proper methods (1), (2). Genome scale metabolic models (GSM) are comprehensive collections of known biochemical reactions, which are catalysed by specific associated proteins coded by genes. GSM method has been successfully used to explain genotype - phenotype relationships for prokaryotic, eukaryotic, unicellular and multi - tissue organisms. GSM has already proven to be successful leading to several important breakthroughs in health and systems medicine fields (3), (4), (5), in biotechnology (6), (7), (8) and many other life science fields (9), (10), (11). GSM is a method for analysing the phenotype responses of an organism by calculating flux distribution and other parameters (12) in order to see carbon distribution potential and environmental perturbation impact on the metabolism (13), (14). Despite the knowledge of genotype in GSM, phenotypic responses are far from full understanding. With advances in omics technologies it has become possible to quantitatively monitor transcripts, proteins and metabolite levels, which makes it possible to narrow this gap between genotype and phenotype (15), (16). In contrast to gene set enrichment analysis, where genes exhibiting similar biological characteristics are sorted and classified in clusters (17), GSM based integration methods also take into account the interconnectivity of a biochemical network, the steady state assumption and Gene - Protein - Reaction (GPR) associations. This allows the analysis of transcriptome data sets to be performed in an interconnected manner on the biochemical network topology (18). There are two fundamental approaches of expression data integration in GSM. Directly integrating transcriptome data into flux bounds (DIRECT) and distributing the reactions into different categories based on transcriptome levels (DISTRIBUTE). The first DIRECT approach was developed by Åkesson (19) they assumed that very low gene expression levels are associated with non-flux reactions. An advanced method E-Flux (20) allows to integrate quantitative transcriptome measurements directly into models as the maximum possible flux value (flux upper bound).

But GIMME (21) algorithm was made to distribute reactions into groups of highly and lowly expressed by comparing them to a user specified threshold.

**IgemRNA** is an open access toolbox for transcriptome data statistical and biochemical network topology-based analysis. IgemRNA was developed in the MATLAB environment in order to take advantage of the up-to-date and most commonly distributed GSM modelling tool Cobra Toolbox 3.0 (22) and spreadsheet file capabilities. Tool is designed to analyse not only the quantitative genome scale transcriptome data measurements like RNA-seq (23) but also a targeted gene group transcriptome data, for example, Gene expression Microarray Analysis (24). IgemRNA allows to analyse transcriptome data directly or to apply GSM biochemical network topology properties and perform optimisation methods like FBA (25) (26) or FVA (27). GSM models without omics data cannot reveal distinct phenotype properties and can predict only the theoretical carbon distribution. By applying transcriptome data to a GSM model, it is possible to explain more precisely an organism’s phenotype properties in different stress and environmental conditions (28). Although many different tools for omics data integration into GSM already exist and have shown to be practical (29), they are not compatible and use a variety of different standards and programming environments (Supplementary Materials 1). This has raised the need in the scientific community for a tool, which combines most of the previously published basic functionalities, can be used in conjunction with already available GSM modelling tools and allows to select a variety of data integration, processing, analysis and storage options in a user-friendly way. IgemRNA has been designed to combine several functionalities of previously published tools (Supplementary Materials 1) like optimisation, transcriptome data initialization and integration methods, and the access is provided via graphical user interface. Additionally, IgemRNA allows the use of medium composition data in combination with transcriptome data from the same environmental conditions. But the main novelty of this tool is the multiple different built - in functions and options for transcriptome data pre-processing including gene mapping and thresholding, and transcriptome non and post GSM based optimisation statistical analysis.

## MATERIALS AND METHODS

IgemRNA is developed in the MATLAB programming environment. All functionality parts of IgemRNA toolbox are open access and free available under the MIT License. GUI window is created with MATLAB *dialog()* function and all user interface controls are added to the window using MATLAB *uicontrol()*. IgemRNA has dependencies for the MATLAB based GSM modelling tool Cobra Toolbox 3.0 (22) functionality and spreadsheet file (xls,xlsx) capabilities to store the results. All IgemRNA’s functionalities are compatible with MATLAB versions not older than 2014 (https://opencobra.github.io/cobratoolbox/latest/installation.html). IgemRNA is available in GITHUB (https://github.com/BigDataInSilicoBiologyGroup/IgemRNA) and is designed for transcriptome data analysis with or without GSM model network topology application.

### Tools architecture description

Cobra Toolbox 3.0 and spreadsheet data files (transcriptomics, GSM model, optional medium data) in xls or xlsx format are required to access all the functionality of IgemRNA (Fig. 1). IgemRNA uses different implemented methods and saves results in spreadsheet files, depending on the user’s selected transcriptome data processing methods and analysis tasks. IgemRNA can also use the built-in methods Flux Balance Analysis (FBA) and Flux Variability Analysis (FVA) of Cobra Toolbox 3.0 (30), to perform more complex analysis on GSM network topology.

**Fig. 1.**
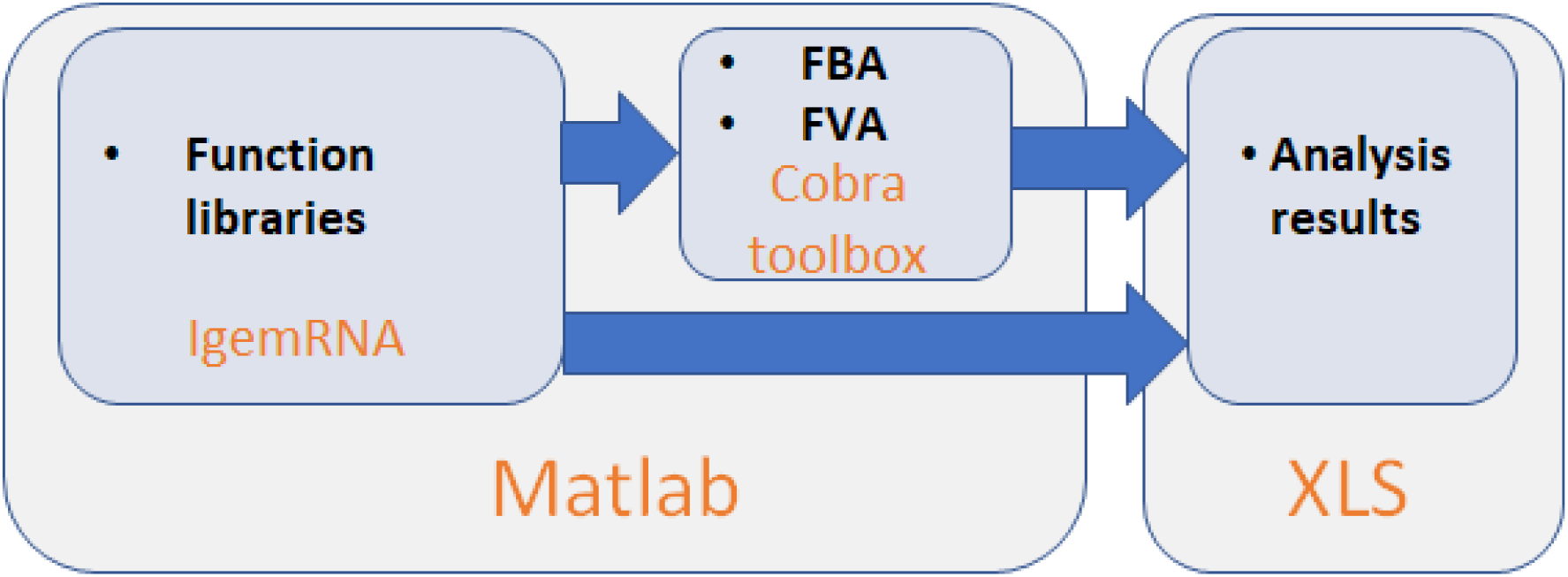
IgemRNA toolbox architecture scheme.

### Tools functionality description

IgemRNA has 7 different functional modules (Fig. 2). IgemRNA modules include the access to the user-friendly interface with data input options, data initialization and pre-processing steps (gene mapping and thresholding), and executes user-selected non and post optimisation analysis tasks. Post optimization tasks utilize some of the Cobra Toolbox 3.0 functions (FBA, FVA). To run IgemRNA tool, user must start the MATLAB software, navigate to the script folder and run *IgemRNA.m*. Depending on the user’s choices in the graphical interface (post-optimization tasks), IgemRNA will launch Cobra Toolbox 3.0. The most time-consuming step is Cobra Toolbox 3.0 initialization with update function. IgemRNA has options to run Cobra Toolbox 3.0 with or without an update function.

**Fig. 2.**
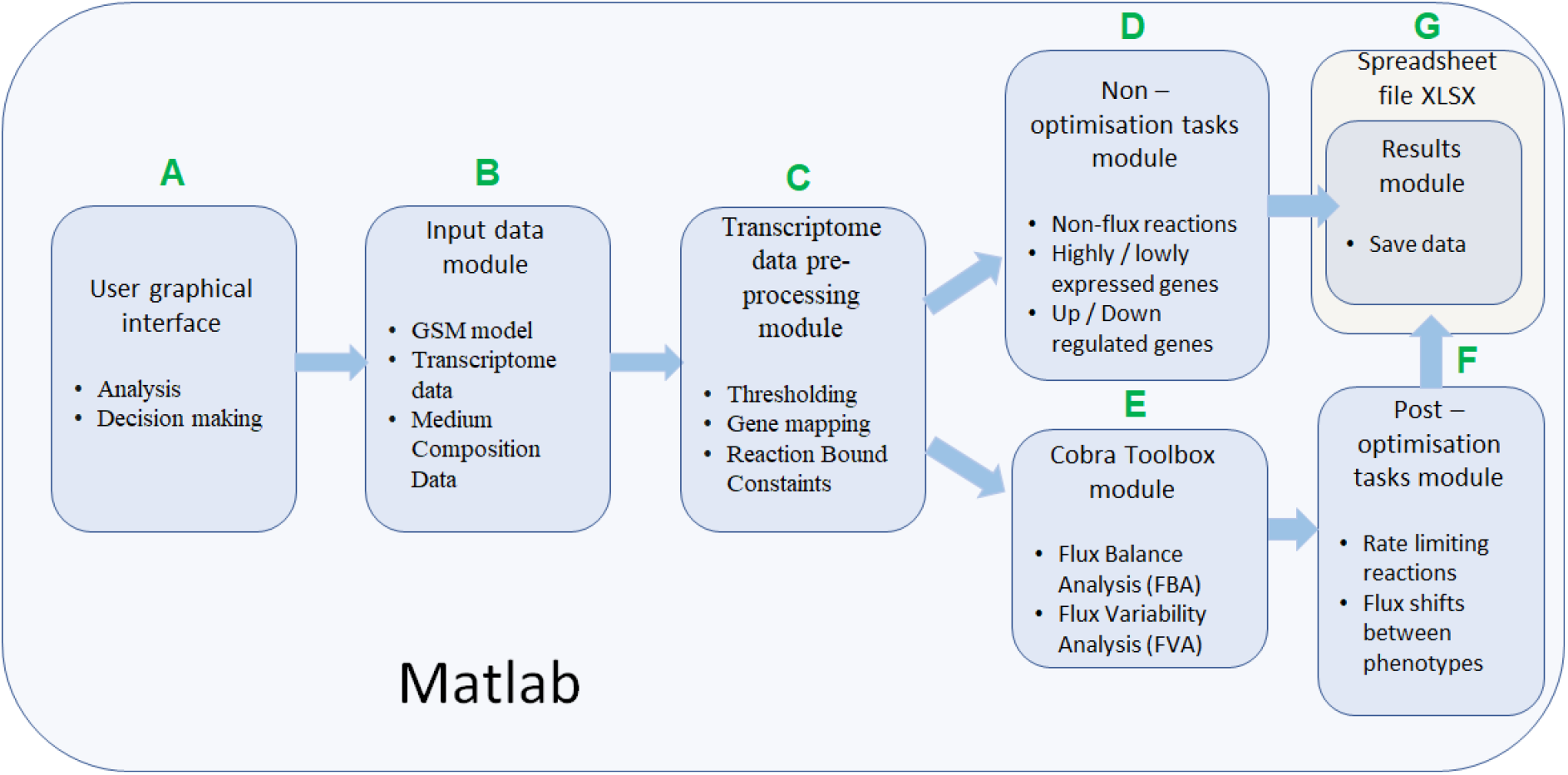
IgemRNA’s function module description

Users can choose which transcriptome data, medium composition and GSM model files to load. Then the user selects desired GPR mapping and thresholding approaches as well as optimization and analysis tasks to perform. When all necessary options have been selected, IgemRNA automatically determines which of non or post optimisation functionalities will be executed. An important functional part of IgemRNA is the **user graphical interface (GUI)** module (Fig. 2A), which collects information about uploaded data files and user’s choices for transcriptome data analysis and optimisation tasks, and passes these parameters to corresponding functional modules. The GUI window is opened using MATLAB *dialog()* function and all user interface controls are added with MATLAB *uicontrol()* function. Depending on what data files the user has supplied certain functional modules are called afterwards. If the user supplies only a transcriptomics data file, optimization task module and post-optimization tasks module will not be executed because they require a GSM model (Fig. 2).

**Input data** module allows to import *transcriptome*, *GSM and medium composition data* from uploaded files (Fig. 2B). All input files must meet specified standards and criteria to be recognised by IgemRNA. Transcriptome data must be located in a spreadsheet XLSX file, where gene names are located in the column with name “GeneId’’ and in the column with name “Data” - the measured transcriptome values. Gene names must be the same as they are defined in the GSM model. Permissible GSM model data file types are SBML, MAT, XLSX and the structure should be the same as defined in Cobra Toolbox 3.0 (22). It is also possible to use BioPax, GPML and SBML files older than 3.0 version before converting them to the newest SBML 3.2 version by online Systems Biology Format Converter (SBFC https://www.ebi.ac.uk/biomodels/tools/converters/). The user can also define uptake rates for substrates. Medium composition file contains data about the medium in which the organism grows, respectively substrate and product reaction rates (mmom*g^-1^ * h^-1^) and optionally a specific growth rate (h^-1^) (Fig. 4B)

**Fig. 3.**
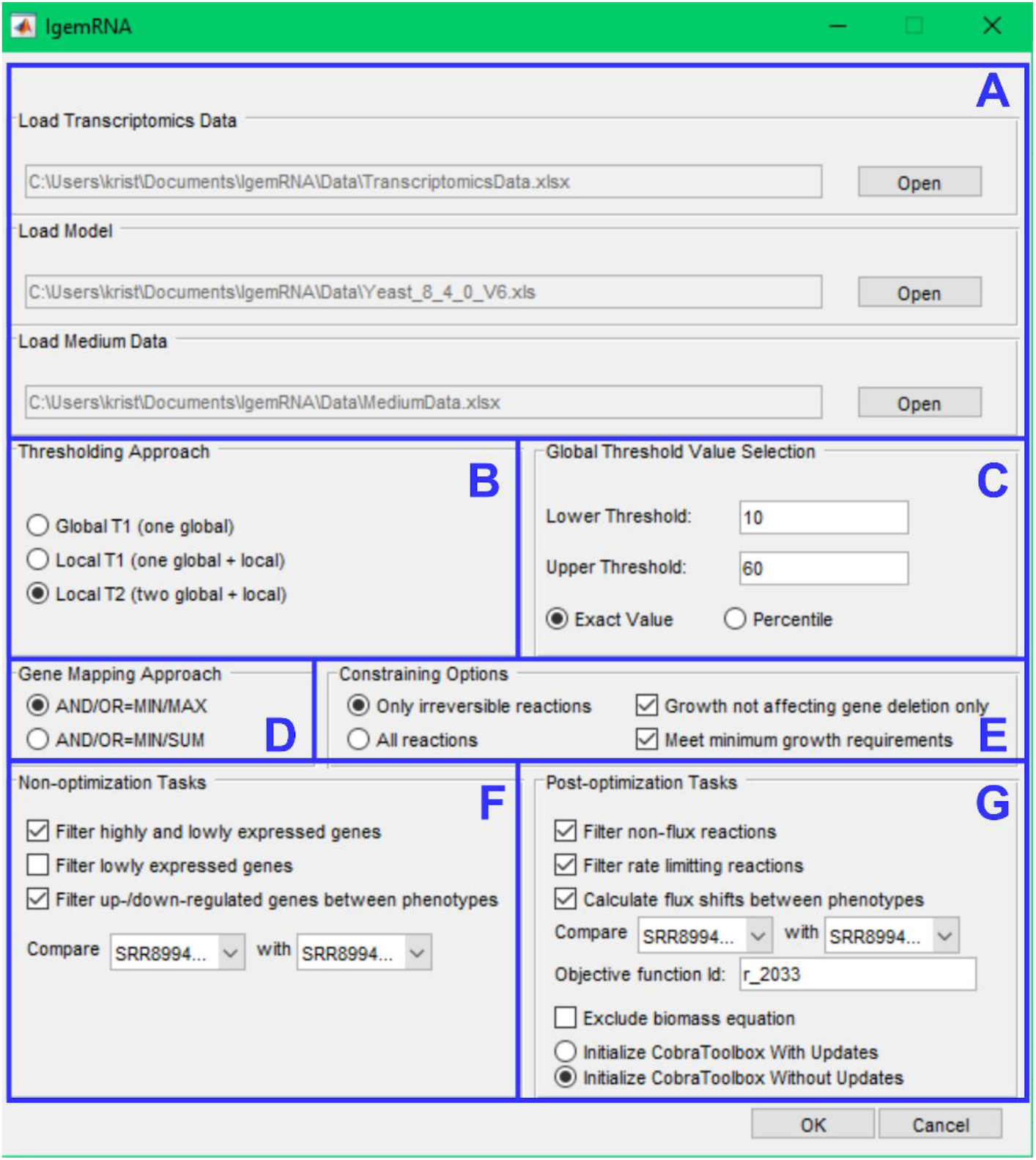
IgemRNA’s graphical interface

**Fig. 4.**
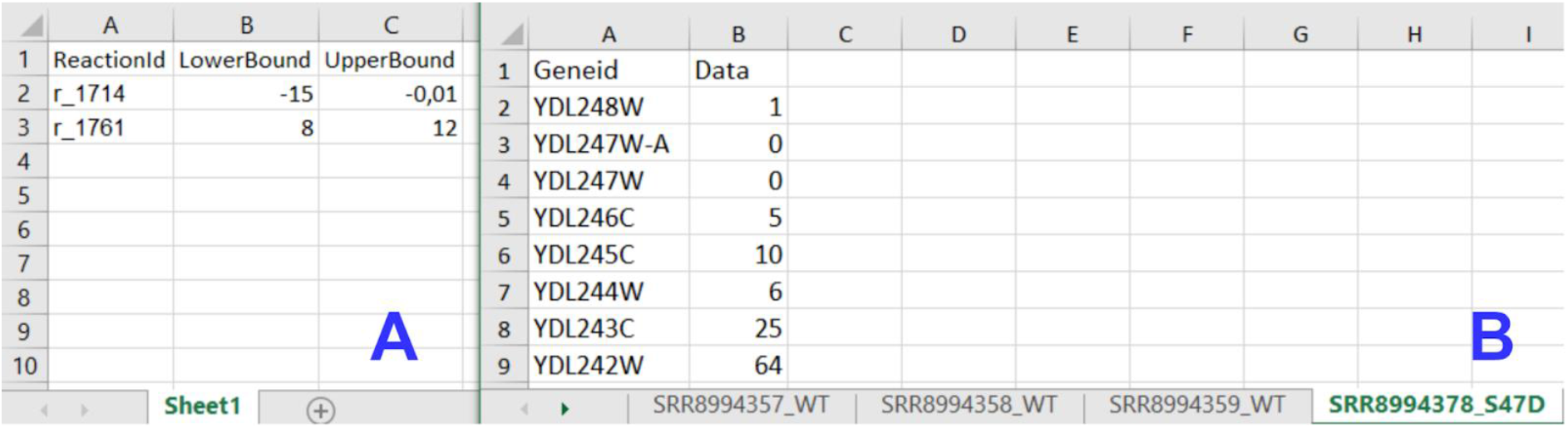
Input data file structure; A - Medium data file structure; B - Transcriptomics data file structure

**Transcriptome pre-processing** functionality module (Fig. 2C) is responsible for preparing transcriptome and GSM model data for analysis and optimisations tasks. Pre-processing of transcriptome data relies mainly on decision parameters: gene mapping and thresholding approach. Different combinations of these decisions influence data processing results and biological data interpretation.

In IgemRNA a **threshold parameter** is required, which defines a border between differentially expressed genes. Two types of thresholds exist: *local* and *global*. *Local* thresholds are automatically calculated for each gene individually given at least two samples of transcriptomics data of the same conditions. *Global* threshold is used as a unique parameter for all genes (31). Different combinations of local and global thresholds lead to different analysis and optimisation results. IgemRNA allows user to choose between 3 different thresholding approaches in data pre-processing step (Table. 1) (https://doi.org/10.1371/journal.pcbi.1007185):

- **Global T1 (GT1)**: is designed to analyse the transcriptome data sets using one global threshold. Example case shows that all transcriptome levels above 130 are considered as expressed and others are considered as suppressed. Global T1 threshold approach can be used for one or several phenotype transcriptome datasets.
- **Local T1 (LT1)**: is designed to analyse transcriptome data sets having one global threshold value and a local rule. Local thresholds are used to set a strict border for specific genes based on their varying gene expression levels across multiple samples in order to determine whether a gene is expressed or suppressed in a specific phenotype case. Local thresholds are only applied to those genes with expression levels above the global threshold since genes with expression levels below the global threshold are automatically seen as suppressed. Example case shows that all transcriptome levels defined by the global threshold above 130 are considered as possibly expressed and below 130 are considered suppressed. Local thresholds for specific genes are used to determine expression or suppression status for genes with expression levels above global threshold.
- **Local T2 (LT2)**: is designed to use 2 different global threshold values: upper threshold and lower threshold. Transcriptome levels higher than the upper global threshold are considered as expressed genes and are active. Transcriptome levels below the lower global threshold are considered as inactive genes. All genes with expression levels between the upper and lower global thresholds are considered as possibly active and local rules for these genes are calculated across multiple gene expression data sets and applied in order to determine their activity levels. The Local T2 thresholding approach can be used if several transcriptome datasets are available. Example of Local T2 shows that all gene expression levels above the upper global threshold 130 are considered as active. Gene expressions lower than global threshold 50 are considered suppressed.

**Table. 1.** Thresholding Options.

| Approach | Threshold input | Visual representation | Example |
| --- | --- | --- | --- |
| **Global T1** | **One global threshold:**<br>exact value input<br>percentile input (25<sup>th</sup>, 75<sup>th</sup>,90<sup>th</sup> ...) | **Global T1** | **Lower Threshold: 130**<br>YDL227C 871<br>YDL226C 126<br>YDL225W 319<br>YDL224C 56<br>YDL223C 13<br>YDL222C 3<br>YDL221W 135 |
| **Local T1** | **One global threshold:**<br>exact value input<br>percentile input (25<sup>th</sup>, 75<sup>th</sup>,90<sup>th</sup> ...)<br>**Multiple local thresholds:**<br>calculated automatically across multiple transcriptomics data samples of the same conditions | **Local T1** | **Lower Threshold: 130**<br>**Local for YDL227C: 600**<br>**Local for YDL225W: 350**<br>YDL227C 871<br>YDL226C 126<br>YDL225W 319<br>YDL224C 56<br>YDL223C 13<br>YDL222C 3<br>YDL221W 135 |
| **Local T2** | **Two global thresholds:**<br>exact value input<br>percentile input (25<sup>th</sup>, 75<sup>th</sup>,90<sup>th</sup> ...)<br>**Multiple local thresholds:**<br>calculated automatically across multiple transcriptomics data samples of the same conditions | **Local T2** | **Lower Threshold: 50**<br>**Upper Threshold: 130**<br>**Local for YDL224C: 60**<br>**Local for YDL226C: 70**<br>YDL227C 871<br>YDL226C 126<br>YDL225W 319<br>YDL224C 56<br>YDL223C 13<br>YDL222C 3<br>YDL221W 135 |

Each threshold approach has unique properties and is included in IgemRNA. Global threshold value input can be manual or automatic. In a manual input scenario user provides an exact expression value whereas in an automatic global threshold input scenario threshold values are calculated based on a user-provided percentile. Local thresholds are always calculated automatically across multiple gene expression data sets during the process of analysis. The selection of thresholding approach is required for both post optimisation and non-optimization cases, where GSM model is not required and is optional.

Other important choice parameters are the **Gene mapping** approach and **Constraining options**. This method is used only for post-optimisation tasks. A GSM model with available GPR data is required. Gene mapping approaches use GPR association, which simplest dogma is one gene - one protein - one reaction in a GSM model. But the presence of enzyme complexes (multiple genes—one protein), isozymes (multiple proteins—one function) and promiscuous enzymes (one protein—multiple functions) in GSM makes these associations more complex and are defined with boolean AND / OR rules. For transcriptome data implementation in GSM one of two approaches of Gene mapping can be applied (Table. 2):

- *Maximum (MIN/MAX)* mapping uses GPR association. AND expression in GPR association is assumed as the smallest gene expression value (MIN), but in OR expression only the highest (MAX) gene expression value is taken.
- *Sum (MIN / SUM)* mapping uses GPR association. AND expression in GPR association is assumed as the smallest gene expression value (MIN), but in OR expression the sum (SUM) of all expression levels is calculated.

**Table. 2.**
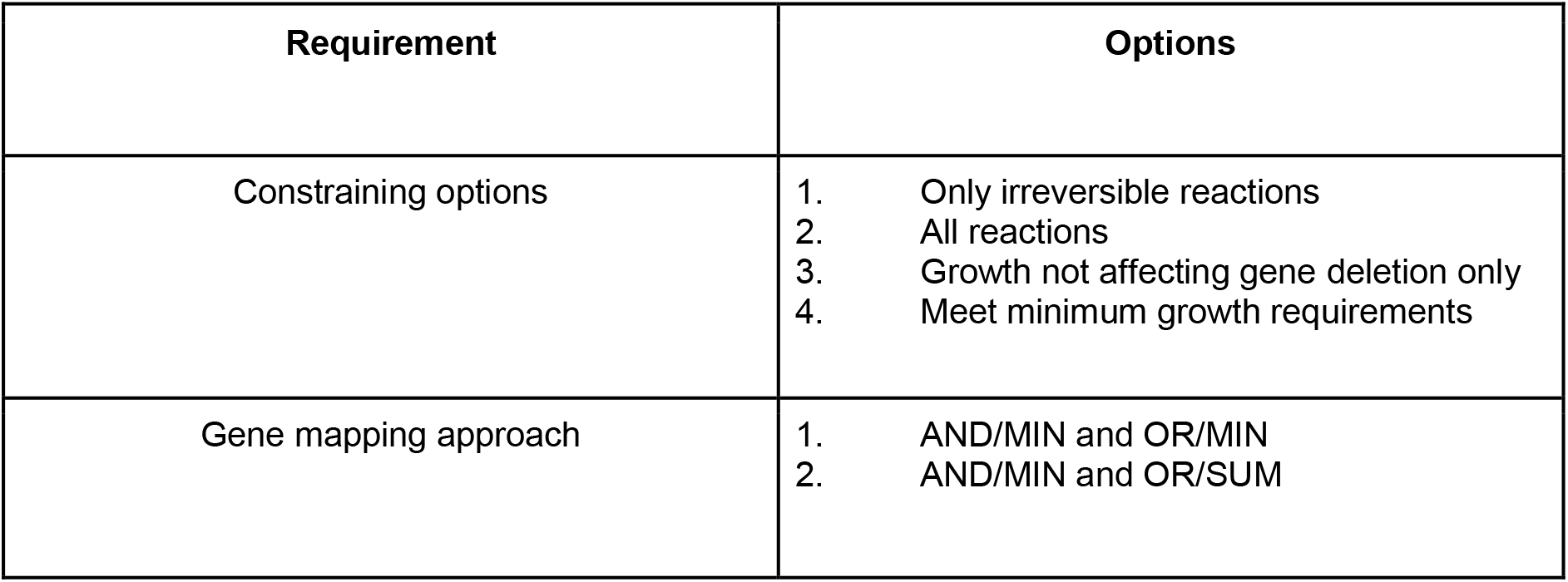
Gene Mapping Options.

| Requirement | Options |
| --- | --- |
| Constraining options | 1. Only irreversible reactions<br>2. All reactions<br>3. Growth not affecting gene deletion only<br>4. Meet minimum growth requirements |
| Gene mapping approach | 1. AND/MIN and OR/MIN<br>2. AND/MIN and OR/SUM |

Cobra Toolbox 3.0 has implemented several previously published transcriptome analysis methods (22). The **constraining options** are used to implement transcriptome absolute values on reaction bounds in the GSM:

- **Only irreversible reactions** function. Enzymatic reactions have 3 different directions in GSM: irreversible, reversible and backward irreversible. This approach constraints only irreversible and backward irreversible reactions in oriented direction.
- **All reactions** function constraints all reactions. Irreversible and backward irreversible reactions in oriented direction, but reversible reactions are constrained in both directions.
- **Growth not affecting gene deletion only** option allows to delete only those genes with expression values below the given threshold and does not affect growth. Cobra Toolbox 3.0 *singleGeneDeletion* analysis with FBA method is performed before executing gene deletion for those genes. Only if the returned output *grRatio* returned by *singleGeneDeletion* function is equal to 1 (meaning that the wild type growth equals to the deletion strain growth) the gene gets deleted.
- **Meet minimum growth requirements** option allows to constrain only those reactions where the gene mapping end value (which is set as a reaction upper bound) is not below the minimum growth requirements for that reaction. Minimum growth requirements are obtained by creating another context-specific model where only the gene deletion and medium exchange reaction constraining is applied in order to calculate the Cobra Toolbox 3.0 FBA (*optimizeCbModel*) minimization of growth.

**Non-optimisation tasks module** is the simplest transcriptome analysis module, which does not require additional Cobra Toolbox 3.0 functionality or a GSM model. This module includes 3 pre-defined different transcriptome data analysis methods:

- Filter highly and lowly expressed genes - method uses chosen threshold data and sorts genes into highly and lowly expressed data sets. Afterwards it is passed to the Results module.
- Filter lowly expressed genes - method uses chosen threshold parameters and filters genes with expression levels below the supplied thresholds and returns them as non-expressed data sets.
- Filter up/down regulated genes between phenotypes - method uses chosen threshold data and filters up and down-regulated genes from two or more transcriptome data sets. The gene names must be the same in all datasets.

All created result data is passed to the Spreadsheet module (Fig. 2G)

**Cobra Toolbox** module is called before post-optimization tasks in order to calculate FBA and FVA results using Cobra Toolbox 3.0 functions (Fig. 2E). This module requires a GSM model.

**Post - optimization task module** is IgemRNA’s advanced transcriptome analysis module which uses GSM metabolic models and analyses them with Cobra Toolbox 3.0. Firstly, the module applies all Transcriptome data pre-processing functions (Fig. 2C). Each function has several optional choices, since each choice in combination with GSM generates different analysis results. This is novelty of IgemRNA in comparison with other tools (supplementary materials 1). Post - optimisation module has several functions for analysing context-specific models:

- **Filter non-flux reactions** - this functionality allows to filter out enzymatic reactions which does not carry a flux, because the coded gene transcription levels are below the chosen threshold value in the Pre-processing module (Fig. 2C).
- **Filter rate limiting reactions** - this functionality finds reaction maximum rates that are equal to the measured transcriptome data levels for genes. Function uses FVA optimisation method to calculate each reaction minimal and maximal rate value and then finds reactions with upper bounds of the same value as the FVA maximal results.
- **Flux shifts between phenotypes** - this function compares minimal and maximal fluxes (calculated by FVA) between different phenotypes or datasets, calculating ratios between them.

All functionalities generate results, which are passed to the **Spreadsheet** module.

Results Module saves all non and post optimization analysis results in spreadsheet files of xls or xlsx format. This is done because further data sorting and statistical analysis is easier to perform in a spreadsheet file.

IgemRNA is available in GITHUB (https://github.com/BigDataInSilicoBiologyGroup/IgemRNA).

### RNA sequencing data analysis

Publicly available and previously published (32) gene expression dataset from *Saccharomyces cerevisiae* BY4741 strain and mutant strain H4-S47D were used for analysis. Reads were aligned using *STAR* aligner (33) and assigned to genomic features using *featureCounts* (34). Statistical comparisons between samples were done using *edgeR* (35). Workflow is publicly available on Galaxy platform (36), (37): https://usegalaxy.eu/u/karlispleiko/w/rna-seq-kp-fromgeosingle-read.

## RESULTS AND DISCUSSIONS

### Available transcriptome data integration tools comparison

Before developing the IgemRNA tool we made a summary of available transcriptome analysis methods. We found many previously published methods and processed them according to various criteria. To classify transcriptome analysis methods, we chose to categorize them by several properties: method name, does return a context-specific model, threshold method, gene mapping approach, requirements to run it, is maintained nowadays, 3rd party software availability and accessibility to build-in statistical analysis methods (Supplementary materials 1).

These methods vary in functionality and data pre-processing approaches. Very first transcriptome analysis method was Åkesson (19), which assumes that the user chosen global threshold of very low gene expression levels is associated with none flux value reactions. GIMME (21) method compares two transcriptome datasets, determines active and inactive genes and minimises lowly expressed reactions thus keeping objective function above set value. iMAT (38) allows the integration of transcriptomic and proteomic data into metabolic models. It groups reactions in highly, moderately and lowly expressed reactions, maximizes highly and minimized lowly expressed reactions. MADE (39) uses two or more sets of expression data across multiple conditions, creates a sequence of binary expression states to find statistically significant changes in gene expression measurements and determines highly/lowly expressed reactions. E-Flux (20) constraints upper bounds for reactions, which are classified as lowly expressed based on a given threshold and expression data. PROM (40) used for integration of transcription regulatory and metabolic networks with gene expression data. Calculates the probability that a gene is active with respect to its transcription factor as specified by expression data and then constrains the reactions maximum flux by a factor of this probability. INIT (41) maximizes reactions based on a qualitative confidence score and minimizes reactions associated with low expression. In addition, this method allows a small net accumulation rate for internal metabolites in order to prevent the removal of necessary reactions. Lee–12 (42) Integrates absolute gene expression data directly into the objective function of a constraint-based model instead of constraining the fluxes. The biological objective function is replaced by a function that minimizes the deviation between gene expression levels and the fluxes. Fang-12 (43) used to predict flux distribution for a perturbed state based on the differences in gene expression levels relative to a reference condition with precalculated flux distribution. This method also allows small variations in the composition of biomass for the perturbed state. RELATCH (44) uses gene expression and fluxomic data from a reference state in order to estimate metabolic changes in a perturbed state for which there is no expression data available. Flux distribution for the perturbed state is calculated by minimizing the adjustment to the reference state. TEAM (45) method estimates time-course flux profiles using temporal gene expression patterns by combining dynamic Flux Balance Analysis (dFBA) and GIMME algorithm. It calculates flux distribution at each time step and uses flux sum minimization in order to find the most optimal solution. GX-FBA (46) method that integrates gene expression data into flux balance analysis, uses the deviation in gene expression levels between a reference state and a perturbed state in order to define flux constraints for the perturbed state.

Akkeson and GIMME, E-Flux uses only one global **threshold**, PROM uses predefined 33rd percentile threshold from average value, INIT uses optional global threshold and positive and negative weights for each reaction, TEAM calculates threshold from M3D microarray dataset database. iMAT uses 2 different global thresholds as lower and upper bounds. Meanwhile, MADE does not use thresholding.

All transcriptome analysis and optimisation tools require a GSM model and one or more transcriptome datasets. Moreover, iMAT requires objective function value, pre-processed transcriptome datasets compatible with reaction bounds. Additionally, to iMAT, GIMME also requires objective function. MADE needs access to a mixed - integer linear program solver (like GUROBI) and more than one transcriptome data set. Therefore, E-Flux requires a function to convert transcriptome data expression levels to fluxes. PROM is developed to use not only transcriptome environmental and genetic perturbation datasets but also regulatory networks for specific cases. Meanwhile RELATCH method requires transcriptome and fluxomics dataset from the same conditions. More advanced method TEAM used initial composition data, temporal transcriptome and biomass composition data.

Along with transcriptome analysis and optimization tools, novelty not less important is tool maintenance, for example Lee-12 has claimed its availability in Cobra Toolbox, but in the newest Cobra Toolbox 3.0 version is not found. Only GIMME, iMAT, INIT and TEAM (only for microarray experiments) are maintained nowadays for our best knowledge.

All published tools are used for only specific cases and are not very informative for other scenarios due to narrow functional potential and a single analysis and optimisation approach. Later published transcriptome analysis approaches are considered ineligible for manuscript classification criteria.

IgemRNA is a novel tool, which has combined many previously listed pre-processing and different transcriptome analysis methods. Tool allows to analyse transcriptome data in conjunction or separately from GSM models to find highly and lowly expressed genes and compare them with other phenotype data sets. IgemRNA has implemented more advanced pre-processing methods than previously listed tools. Thresholding approaches include 3 different options, gene mapping has 4 different approaches.

IgemRNA is compatible with MATLAB based software and optionally uses Cobra Toolbox 3.0 functionality. In comparison with previously mentioned methods, IgemRNA facilitates several different thresholding, gene mapping approaches and constraining options for transcriptome data integration into GSM models. Additional feature is the possibility for the end user to select what kinds of data to extract from the result context-specific models. More advanced analysis methods in IgemRNA also include data comparison between different phenotypes. On top of that, all previously mentioned options can easily be selected via graphical user interface.

### IgemRNA demonstration

The main novelty within this paper is the IgemRNA tool itself, combining multiple different key decisions and options for transcriptomics data analysis and integration in GSM models. IgemRNA’s graphical interface alleviates the complexity of data input and pre-processing (thresholding, gene mapping) for the end user (Fig. 3). Another useful feature of IgemRNA is the possibility for the user to choose which of different data analysis tasks to perform including both non-optimization and post-optimization tasks.

After running the main IgemRNA’s script *IgemRNA.m*, a dialog box is opened where the user must first select the transcriptomics data file of xls or xlsx format in the file upload section (Fig. 3A). The transcriptomics data should be organized into two columns ‘GeneId’ and ‘Data’ (Fig. 4B) where ‘GeneId’ corresponds to the genes in the GSM model if one is supplied. The transcriptomics data file can consist of multiple RNA-seq samples of the same conditions as well as different phenotypes. Therefore, the phenotype and sample name must appear as the sheet name since this name will be used for naming the result files and selecting phenotypes for comparison (FIG. 3F, G).

Having selected a transcriptomics data file, the non-optimization tasks section (Fig. 3F) becomes visible and having selected a GMS model of Cobra Toolbox 3.0 available formats, the post-optimization tasks section (Fig. 3G) becomes visible. Optionally, the user can also supply a medium data file of xls or xlsx format in the file upload section (Fig. 3A) where all the needed reactions as well as their upper and lower bounds are listed (Fig. 4A).

Next 4 dialog sections (Fig. 3 B, C, D, E) contain options for data pre-processing. **Thresholding approach** (Fig. 3B) and **global threshold value selection** (Fig. 3C) options determine methods for transcriptomics data pre-processing in order to split genes into groups of highly and lowly expressed. It is possible to choose one of three thresholding approaches: Global T1 (GT1), Local T1 (LT1) or Local T2 (LT2) (see 2.2 threshold parameter). Depending on the selected approach, the user is then asked to enter one or two global threshold values which can be supplied as a percentile or an exact value. **Gene mapping approach** (Fig. 3D) and **constraining options** (Fig. 3E) determine methods for transcriptomics data integration in a GSM model. Gene mapping approach specifies what operations to use for mapping gene expression data to gene-protein-reaction (GPR) associations (see 2.2 Gene mapping) whereas constraining options allow to constrain only irreversible or all reactions. In addition, it is possible to constrain reactions in a way that preserves growth (see 2.2 Constraining options) by deleting only those genes that do not affect growth (Growth not affecting gene deletion only) and constraining only those reactions where the gene mapping end value is not below the minimum requirements for growth (Meet minimum growth requirements).

**Non-optimization tasks** (Fig. 3F) contain several options for transcriptomics data analysis: filtering highly and lowly expressed genes and saving results in excel, filtering only lowly expressed genes and saving results in excel and comparing gene expression data between phenotypes. **Post-optimization tasks** combine transcriptomics data and a GSM model in order to extract data on reaction level. It is possible to filter non-flux reactions, rate limiting reactions and calculate reaction flux shifts between different phenotypes. An objective function is required in order to perform optimizations on the GSM model and since post-optimization tasks use some of the Cobra Toolbox 3.0 functions, the user is also asked to choose how to start Cobra Toolbox: with or without updates.

To validate IgemRNA we used *Sacharomyces cerevisiae* consensus genome scale metabolic model (47) version 8.4.0 (48) and appropriate transcriptome validation dataset was taken from (32), where were performed high-throughput genetic screenings that provide a novel global map of the histone residues required for transcriptional reprogramming in response to heat and osmotic stress in steady state growth conditions. All validation details are found in supplementary material 2.

In order to perform test cases provided in this user manual, simply run the provided test case scripts via MATLAB environment having initialized CobraToolbox 3.0 beforehand. Test case script file names are given at the end of each test case section, for example *TestCase_determineGeneActivity.m* script will run non-optimisation task filter highly and lowly expressed genes in supplementary material 2.

## CONCLUSIONS

Having summarized several transcriptome data processing tools, we found that many of them are case specific and allow users to select only some of the possible data pre-processing and analysis functions. More recently proposed tools however are not as widely used due to the complexity of required input data, for example, tools that facilitate time series data or genome-scale metabolomics data which are not widely accessible or use their own unique FBA modified functions or different 3rd party software.

IgemRNA novelty is the possibility of applying several different thresholding, gene mapping approaches and constraining options for transcriptome data integration into GSM models as well as the feature of allowing the end user to select what kinds of data to extract from the result context-specific models.

IgemRNA in contrast to Gene set enrichment analysis additionally validates transcriptomics measurements quality, where minimal metabolic network connectivity and flux requirements must be fulfilled, otherwise transcriptome data quality is questionable.

IgemRNA has a user-friendly graphical interface and facilitates Cobra Toolbox 3.0 standards for optimisation tasks, which does not require additional user skills.

## DATA AVAILABILITY

IgemRNA MATLAB scripts and test cases for transcriptome data integration in genome scale metabolic models are available at https://github.com/BigDataInSilicoBiologyGroup/IgemRNA

## SUPPLEMENTARY DATA

All IgemRNA test data and scripts are found at https://github.com/BigDataInSilicoBiologyGroup/IgemRNA and are available at NAR online.

## CONFLICT OF INTEREST

### No conflict of interest exists

We wish to confirm that there are no known conflicts of interest associated with this publication and there has been no significant financial support for this work that could have influenced its outcome.

## Author contributions

**Kristina Grausa**: Methodology, Software, Data curation, Investigation, Validation, Writing- Reviewing and Editing. **Agris Pentjuss**: Conceptualization, Methodology, Investigation, Supervision, Writing- Original draft preparation, Script validation. **Karlis Pleiko**: RNA sequencing methodology, Writing, **Ivars Mozga**: Data curation, Writing.

