## Supplementary data for "Integrative Gene Expression and Metabolic Analysis tool IgemRNA"

### Supplementary Material 1 - Comparison of methods for transcription data integration

| Name | Year | Returns a context - specific model | Thresholding | Gene mapping | Data Requirements | Is maintained nowadays | Availability in third party software |
|---|---|---|---|---|---|---|---|
| Akesson | 2003 | No | One user-specified threshold | Not found | A genome-scale metabolic network reconstruction. | No | Not found |
| Gene Inactivity Moderated by Metabolism and Expression (GIMME) | 2008 | Yes | One user-specified threshold | Expression data has to be mapped to reactions before calling the algorithm, can be done using Cobra Toolbox pre-processing functions | 1. A genome-scale metabolic network reconstruction,<br>2. Specified objective function,<br>3. one threshold,<br>4. one gene expression data set,<br>5. calculated reaction expression levels (gene mapping) | Yes | MATLAB,<br>Cobra Toolbox |
| Integrative metabolic analysis tool (iMAT) | 2008 | Yes | Two user-specified thresholds: lower threshold and upper threshold | | 1. A genome-scale metabolic network reconstruction,<br>2. two specified thresholds,<br>3. one expression data set,<br>4. calculated reaction expression levels (gene mapping) | Yes | MATLAB,<br>Cobra Toolbox |
| Metabolic Adjustment by Differential Expression (MADE) | 2011 | Returns Flux distribution | No need for a user-specified threshold | Not found | 1. A genome-scale metabolic network reconstruction,<br>2. two or more gene expression data sets<br>3. a mixed-integer linear program solver | No | Linux / Windows/ MacOsX<br>MATLAB (requires a mixed-integer linear program solver)<br><br>(The most recent version of MADE, |

| | | | | | | | |
|---|---|---|---|---|---|---|---|
| | | | | | | | along with other tools for integrating expression data, is available as part of the TIGER software package.) |
| E-Flux | 2009 | Returns Flux distribution | One user-specified threshold | Not found | 1. A genome-scale metabolic network reconstruction,<br>2. a function to convert expression levels into an upper bound on fluxes,<br>3. gene expression data,<br>4. threshold | Not found | Not found |
| Probabilistic regulation of metabolism (PROM) | 2010 | Returns Flux distribution | A predefined low threshold (default: 33rd percentile) | Not found | 1. A genome-scale metabolic network reconstruction,<br>2. a range of expression data from various environmental and genetic perturbations,<br>3. a regulatory network structure | Not found | Not found |
| Integrative Network Inference for Tissues algorithm (INIT) | 2012 | Yes | Optional minimum flux threshold for expressed reactions (default 1e-8) and positive or negative weights for each reaction | Expression data has to be mapped to reactions before calling the algorithm | 1. A genome-scale metabolic network reconstruction,<br>2. gene expression data | Yes | MATLAB,<br>Cobra Toolbox |
| Lee–12 | 2012 | Returns Flux distribution | Not found | Not found | 1. A genome-scale metabolic network reconstruction,<br>2. Gene expression data | Not found | Cobra Toolbox (MATLAB) |
| Fang–12 | 2012 | Returns Flux distribution | Not found | Not found | Not found | Not found | Not found |

| | | | | | | | |
|---|---|---|---|---|---|---|---|
| RELATive CHange (RELATCH) | 2012 | Returns Flux distribution | Not found | Not found | 1. gene expression data and fluxomic data from a reference state | Not found | Not found |
| Temporal Expression-based Analysis of Metabolism (TEAM) | 2012 | Returns Flux distribution | Threshold determination using background M3D data set | Not found | 1. Initial media composition data<br>2. Temporal gene expression data<br>3. Temporal biomass data | Accessible but only for microarray experiments | Not found |
| Gene-expression FBA (GX–FBA) | 2012 | Returns Flux distribution | Not found | Not found | Not found | Not found | Not found |
| IgemRNA | 2021 | Yes | 1. Global T1 (GT1)<br>2. Local T1 (LT1)<br>3. Local T2 (LT2) | Included, possible options:<br>1. Only irreversible reactions,<br>2. All reactions,<br>3. Growth not affecting gene deletion only,<br>4. Meet minimum growth requirements,<br>5. AND/OR = MIN/MAX,<br>6. AND/OR = MIN/SUM | 1. A genome-scale metabolic network reconstruction,<br>2. gene expression data,<br>3. external metabolite uptake data (optional),<br>4. threshold/-s | Yes | MATLAB, Cobra Toolbox |

#### Supplementary Material 2. IgemRNA test case scenario

IgemRNA is a library with a graphical user interface written for the MATLAB environment and facilitates some of the Cobra Toolbox 3.0 functionality. IgemRNA performs not only Gene sets enrichment analysis-based functions, but also allows integrate transcriptomics data in metabolic models. Also, IgemRNA allows validate transcriptomics data facilitating interconnectivity of biochemical networks, steady state assumptions, Gene - Protein - Reaction relationship and can use optional medium composition data to create context-specific models.

##### Folder structure description

Files are extracted from the archive (https://github.com/BigDataInSilicoBiologyGroup/IgemRNA). The IgemRNA tool consists of four root folders (Data, Scripts and Results non-optimization, Results post-optimization) and an *IgemRNA.m* file which calls the user graphical interface form. Data folder is where the input data files are stored, initially this folder contains the data files used for this demonstration:

a. *MediumData.xlsx* (medium composition data)

b. *Yeast_8_4_0.xls* (the yeast consensus genome-scale model)

c. *TranscriptomicsData.xlsx* (RNA-req measurements) (available https://www.ncbi.nlm.nih.gov/geo/query/acc.cgi?acc=GSE130549)

Transcriptomics data and medium composition data can be provided as an .xls or an .xlsx file and must meet the following format (Fig. 1) where shown columns are provided and named accordingly and sheet names correspond to a phenotype name see IgemRNA demonstration section in main manuscript.

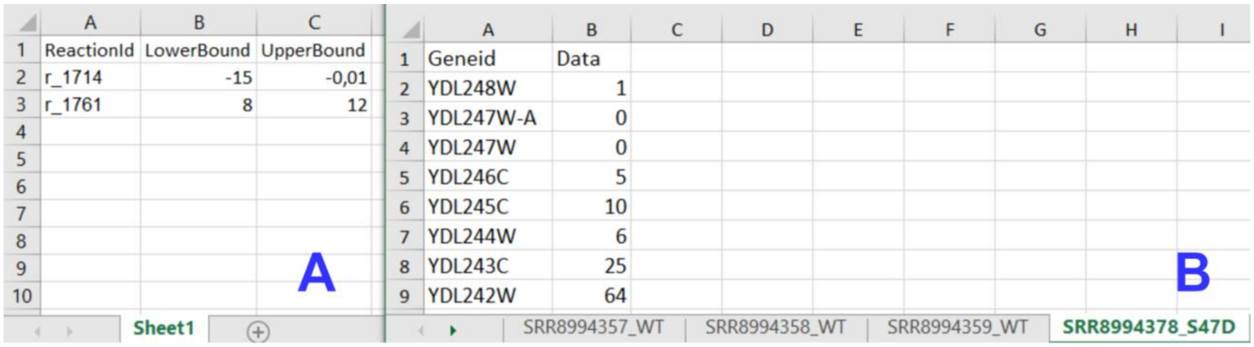

| | A | B | C |
|---|---|---|---|
| 1 | ReactionId | LowerBound | UpperBound |
| 2 | r_1714 | -15 | -0,01 |
| 3 | r_1761 | 8 | 12 |



| | A | B |
|---|---|---|
| 1 | Geneid | Data |
| 2 | YDL248W | 1 |
| 3 | YDL247W-A | 0 |
| 4 | YDL247W | 0 |
| 5 | YDL246C | 5 |
| 6 | YDL245C | 10 |
| 7 | YDL244W | 6 |
| 8 | YDL243C | 25 |
| 9 | YDL242W | 64 |



Fig. 1. Input data file structure; A - Medium data file structure;

B - Transcriptomics data file structure

The model can be provided in xls, sbml or other formats supported by Cobra Toolbox 3.0.

Scripts folder consists of all the script files that are being executed by the IgemRNA tool according to user's selections in the IgemRNA form as well as the test cases provided in this demonstration. The Results non-optimization and Results post-optimization folders are where all the result files are being saved. These folders are initially empty (for more details see section in main publication Materials and Methods Tools functionality description).

#### Starting IgemRNA tool

In order to start the IgemRNA tool a user must start the Matlab environment and run the *IgemRNA.m* script located in the root folder. This script opens the graphical user interface of IgemRNA (Fig. 2).

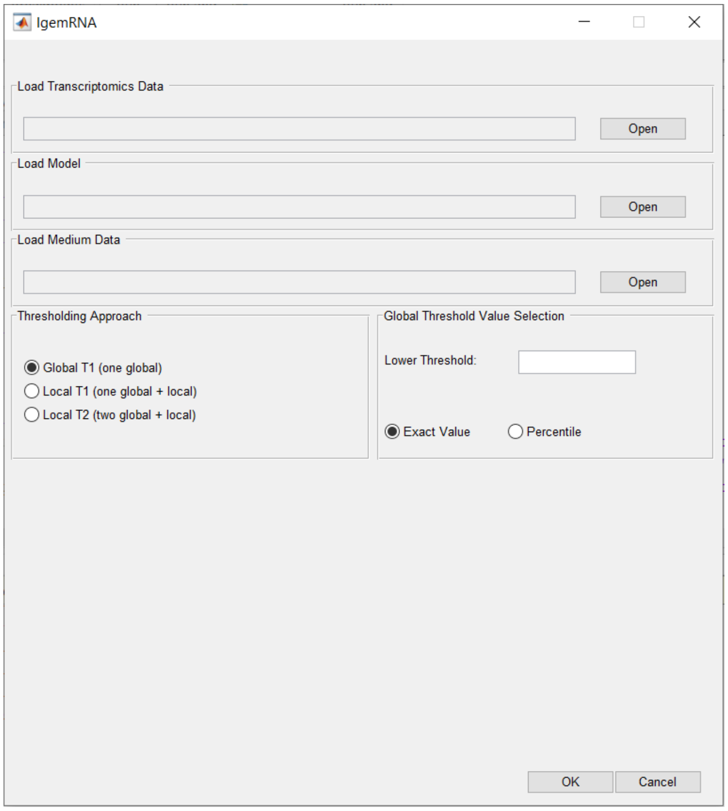


Fig. 2. IgemRNA start window

#### File upload

To access all options in the IgemRNA form, the user must supply input data files Fig. 3 A section. This can be done by pressing the 'Open' button in the corresponding file row and finding the necessary files via File Explorer. Transcriptomics data is required to run non-optimization tasks (Fig. 3 F) but an additional model file is necessary to access the post-optimization tasks (Fig. 3 G). Medium composition file is optional if such data is available, the selection of this data file does not extend the form, but specifies the given exchange reaction constraints (upper bounds and lower bounds) on the model for post-optimization tasks analysis. For an organized overview of the analysis and results it is recommended that the necessary data files are located in the Data folder of IgenRNA tool (for more details see main publication section Materials and Methods Tools functionality description).

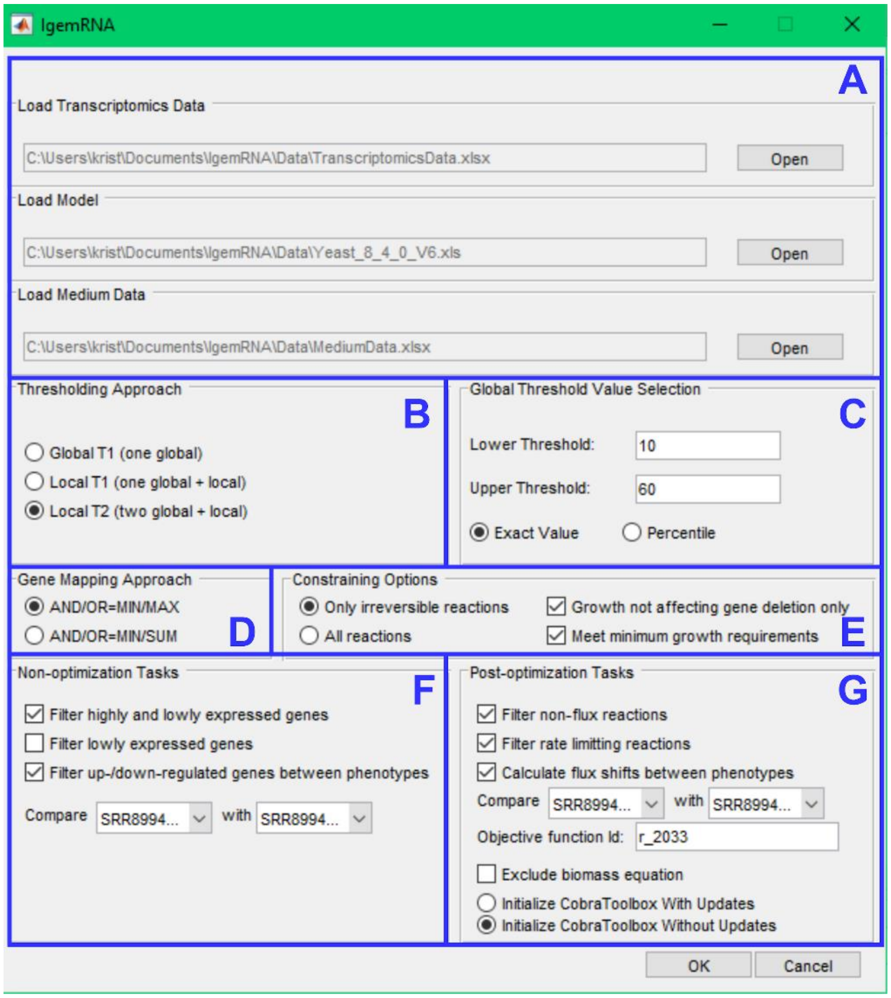


Fig. 3. Full IgemRNA form

**Running test cases**

In order to perform test cases provided in this user manual, simply run the provided test case scripts via MATLAB environment having initialized CobraToolbox 3.0 beforehand. Test case script file names are given at the end of each test case section.

**Non-optimization tasks**

Non-optimization tasks include several transcriptomics data analysis tasks: filter highly and lowly expressed genes, filter lowly expressed genes, filter up/down regulated genes between different phenotypes or data sets. The results for each task are stored in a different folder within the 'Results non-optimization' folder: Gene expression level comparison, Highly-lowly expressed genes, Lowly expressed genes (Fig. 4).

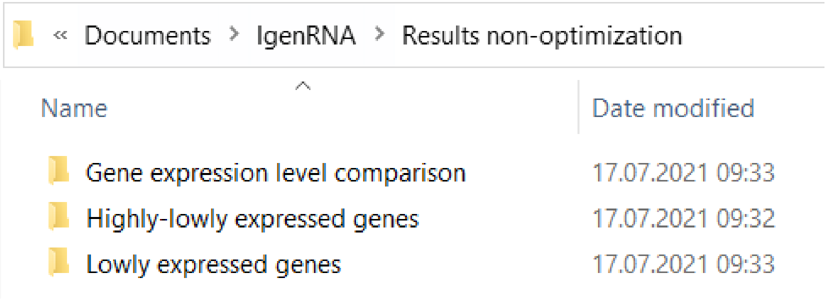


Fig. 4**.** Non-optimization results folder

**a. Filter highly and lowly expressed genes**

Non-optimization task 'Filter highly and lowly expressed genes' generates result excel files for each provided transcriptomics data set. File names are assigned based on the provided dataset and phenotype name (from transcriptomics data), the selected thresholding approach (GT1, LT1, LT2) and provided global thresholds values (Fig. 5).

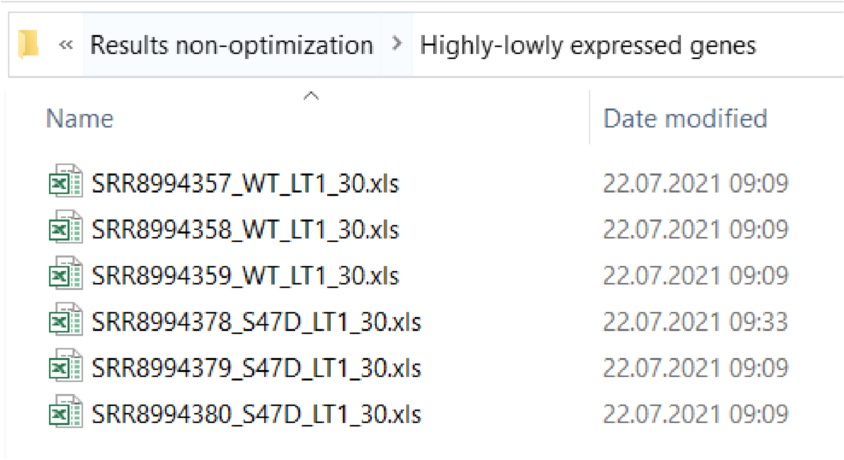


Fig. 5. Highly-lowly expressed genes folder

Each excel file contains one sheet with the list of genes provided by transcriptomics data and 4 columns: GeneId, Data (the expression value), ExpressionLevel and LocalThApplied. The ExpressionLevel column contains the expression levels determined based on the chosen thresholding approach, provided global and for thresholding approaches (LT1 and LT2) calculated local thresholds. Column LocalThApplied displays whether a local threshold for a specific gene was applied (Fig. 6).

| | A | B | C | D |
|---|---|---|---|---|
| 1 | GeneId | Data | ExpressionLevel | LocalThApplied |
| 2 | YDL248W | 1 | low | 0 |
| 3 | YDL247W-A | 0 | low | 0 |
| 4 | YDL247W | 0 | low | 0 |
| 5 | YDL246C | 5 | low | 0 |
| 6 | YDL245C | 10 | low | 0 |
| 7 | YDL244W | 6 | low | 0 |
| 8 | YDL243C | 25 | low | 1 |
| 9 | YDL242W | 64 | high | 0 |
| 10 | YDL241W | 20 | low | 1 |
| 11 | YDL240C-A | 1 | low | 0 |
| 12 | YDL240W | 9 | low | 0 |
| 13 | YDL239C | 0 | low | 0 |
| 14 | YDL238C | 5 | low | 0 |
| 15 | YDL237W | 13 | low | 0 |
| 16 | YDL236W | 10 | low | 0 |
| 17 | YDL235C | 12 | low | 0 |
| 18 | YDL234C | 134 | high | 0 |
| [illegible] | [illegible] | [illegible] | [illegible] | [illegible] |

Fig. 6. Filter highly and lowly expressed genes result file (thresholding approach LT1)

To perform this test case run the file *TestCase_determineGeneActivity.m* in the ‘Scripts’ folder of IgemRNA tool.

##### b. Filter lowly expressed genes

Non-optimization task ‘Filter lowly expressed genes’ generates separate excel result files for each dataset provided in transcriptomics data file. These result files contain filtered gene lists including genes that are below a given threshold based on the selected thresholding approach. File names include dataset and phenotype name (from transcriptomics data file), thresholding approach (GT1, LT1, LT2) name and provided global threshold values (Fig. 7).

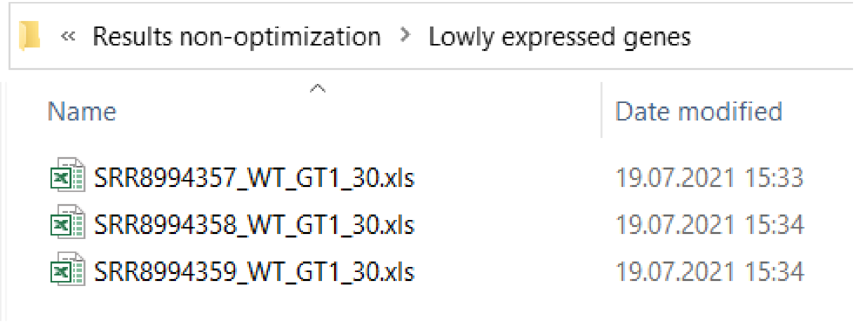


Fig. 7. Non-optimization results folder

The result for lowly expressed genes coincides with the provided transcriptomics data format. Each file consists of two columns, gene identificator and the expression level, but only those genes that are below a given threshold (depending on which thresholding approach is applied) are listed in the result files. The test case provided for this task shows genes with expression level below 30 using the Global T1 approach (Fig. 8).

| | A | B |
|---|---|---|
| 1 | YDL248W | 0 |
| 2 | YDL247W-A | 1 |
| 3 | YDL247W | 0 |
| 4 | YDL246C | 0 |
| 5 | YDL245C | 14 |
| 6 | YDL244W | 3 |
| 7 | YDL243C | 3 |
| 8 | YDL241W | 13 |
| 9 | YDL240C-A | 1 |
| 10 | YDL240W | 0 |
| 11 | YDL239C | 2 |
| 12 | YDL238C | 0 |
| 13 | YDL237W | 11 |
| 14 | YDL236W | 14 |
| 15 | YDL235C | 6 |
| 16 | YDL234C | 13 |
| 17 | YDL233W | 3 |

Sheet1

Fig. 8. Lowly expressed genes result file

To perform this test case run the file *TestCase_filterLowlyExpressedGenes.m* in the 'Scripts' folder of IgemRNA tool.

**c. Filter up/down regulated genes between phenotypes**

Non-optimization task 'Filter up/down regulated genes between phenotypes' generates result excel files in the 'Gene expression level comparison' folder. Result file names contain dataset and phenotype names for both transcriptomics datasets that have been compared (Fig. 9).

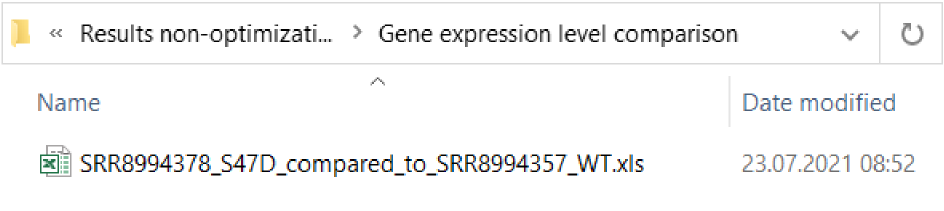


Fig. 9. Up/Down regulated genes in comparison to another phenotype

These results excel data files contain a full gene list from the target dataset and the corresponding genes that are found in the source dataset (Fig. 10. A column). Expression values for both target and source dataset are displayed (Fig. 10. B, C columns) as well as the determined up/down regulation status (Fig. 10. D column).

| | A | B | C | D |
|---|---|---|---|---|
| 1 | GeneId | SRR8994378_S47D_ExpressionValue(target) | SRR8994357_WT_ExpressionValue(source) | Up/Down regulated |
| 2 | YDL248W | 1 | 0 | up |
| 3 | YDL247W-A | 0 | 1 | down |
| 4 | YDL247W | 0 | 0 | equal |
| 5 | YDL246C | 5 | 0 | up |
| 6 | YDL245C | 10 | 14 | down |
| 7 | YDL244W | 6 | 3 | up |
| 8 | YDL243C | 25 | 3 | up |
| 9 | YDL242W | 64 | 48 | up |
| 10 | YDL241W | 20 | 13 | up |
| 11 | YDL240C-A | 1 | 1 | equal |
| 12 | YDL240W | 9 | 0 | up |
| 13 | YDL239C | 0 | 2 | down |
| 14 | YDL238C | 5 | 0 | up |
| 15 | YDL237W | 13 | 11 | up |
| 16 | YDL236W | 10 | 14 | down |
| 17 | YDL235C | 12 | 6 | up |
| 18 | YDL234C | 134 | 13 | up |

Fig. 10. Up/Down regulated genes in comparison to another phenotype result file

To perform this test case run the file *TestCase_compareGeneExpressionLevels.m* in the 'Scripts' folder of IgemRNA tool.

**Post-optimization tasks**

Context-specific models generated by IgemRNA post-optimization tasks as well as the results of the analysis performed on these models are saved in the 'Results post-optimization' folder of IgemRNA tool (Fig. 11). The post-optimization tasks are saved in the folders with the corresponding name: Flux-shifts, Non-flux reactions and Rate limiting reactions (for more details see section Materials and Methods Tools functionality description).

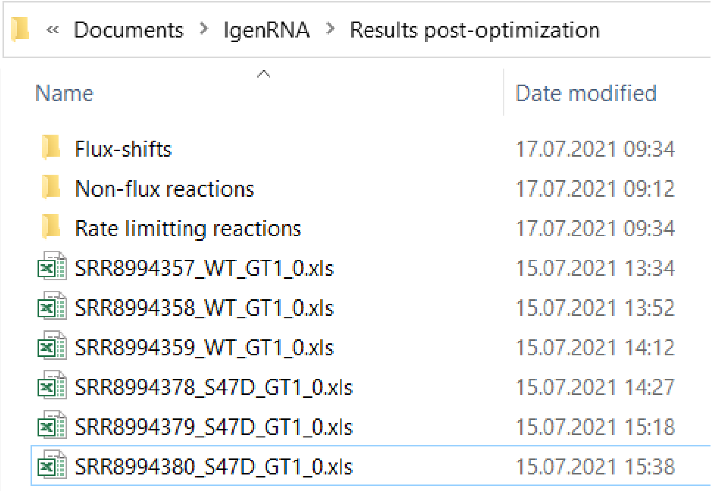


Fig. 11. Results folder after post-optimization task execution

To generate context-specific models used for these test cases run the file *TestCase_generateContextSpecificModels.m* in the 'Scripts' folder of IgemRNA tool. Since this script takes a long time to execute, the context-specific model files used for this demonstration have already been provided in the 'Results post-optimization' folder.

**a. Filter non-flux reactions**

Post-optimization task 'filter non-flux reactions' performs an analysis on the created context-specific models of the same phenotype, the name of the phenotype is included in the result file name (Fig. 12). This analysis filters those reactions that carry no flux.

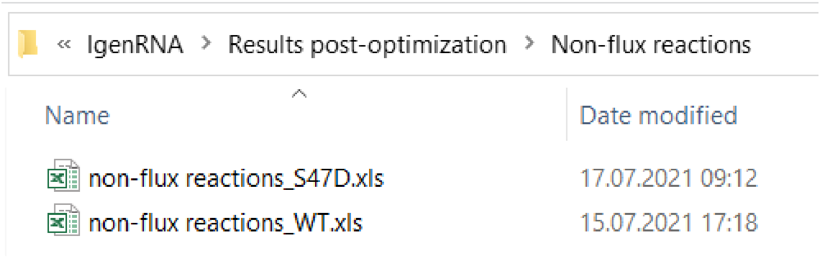


Fig. 12. Non-flux reactions result folder

Each result excel file contains a list for each transcriptomics dataset of reactions that carry no flux in the result context-specific model created by integration of the supplied transcriptomics data into the provided model. An additional sheet for all the common non-flux reactions of the same phenotype is also provided in the sheet 'Common_(phenotype name)' (Fig. 13).

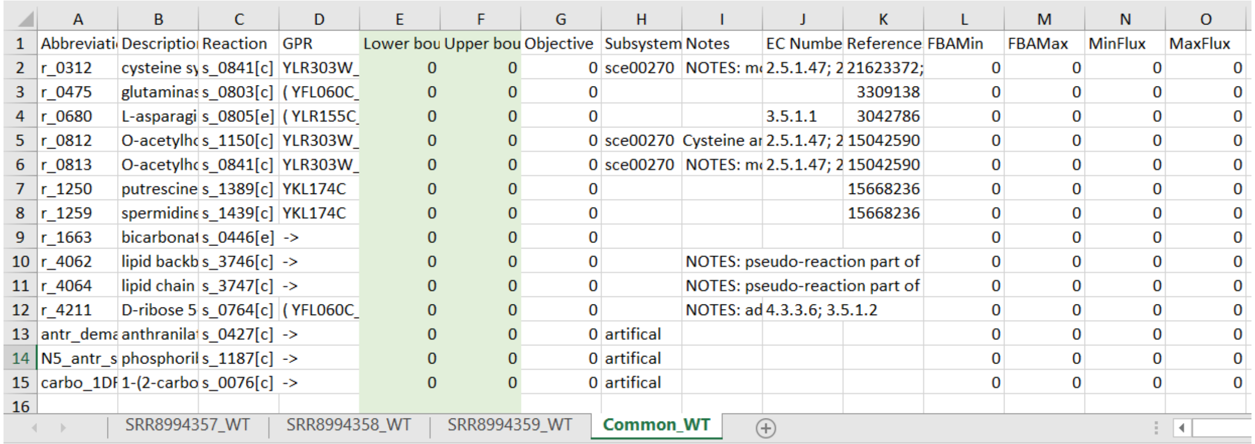

| | A | B | C | D | E | F | G | H | I | J | K | L | M | N | O |
|---|---|---|---|---|---|---|---|---|---|---|---|---|---|---|---|
| 1 | Abbreviati | Descriptio | Reaction | GPR | Lower bou | Upper bou | Objective | Subsystem | Notes | EC Numbe | Reference | FBAMin | FBAMax | MinFlux | MaxFlux |
| 2 | r_0312 | cysteine sy | s_0841[c] | YLR303W_ | 0 | 0 | 0 | sce00270 | NOTES: m | 2.5.1.47; 2 | 21623372; | 0 | 0 | 0 | 0 |
| 3 | r_0475 | glutaminas | s_0803[c] | ( YFL060C_ | 0 | 0 | 0 | | | | 3309138 | 0 | 0 | 0 | 0 |
| 4 | r_0680 | L-asparagi | s_0805[e] | ( YLR155C_ | 0 | 0 | 0 | | | 3.5.1.1 | 3042786 | 0 | 0 | 0 | 0 |
| 5 | r_0812 | O-acetylhc | s_1150[c] | YLR303W_ | 0 | 0 | 0 | sce00270 | Cysteine ar | 2.5.1.47; 2 | 15042590 | 0 | 0 | 0 | 0 |
| 6 | r_0813 | O-acetylhc | s_0841[c] | YLR303W_ | 0 | 0 | 0 | sce00270 | NOTES: m | 2.5.1.47; 2 | 15042590 | 0 | 0 | 0 | 0 |
| 7 | r_1250 | putrescine | s_1389[c] | YKL174C | 0 | 0 | 0 | | | | 15668236 | 0 | 0 | 0 | 0 |
| 8 | r_1259 | spermidine | s_1439[c] | YKL174C | 0 | 0 | 0 | | | | 15668236 | 0 | 0 | 0 | 0 |
| 9 | r_1663 | bicarbonat | s_0446[e] | -> | 0 | 0 | 0 | | | | | 0 | 0 | 0 | 0 |
| 10 | r_4062 | lipid backb | s_3746[c] | -> | 0 | 0 | 0 | | NOTES: pseudo-reaction part of | | | 0 | 0 | 0 | 0 |
| 11 | r_4064 | lipid chain | s_3747[c] | -> | 0 | 0 | 0 | | NOTES: pseudo-reaction part of | | | 0 | 0 | 0 | 0 |
| 12 | r_4211 | D-ribose 5 | s_0764[c] | ( YFL060C_ | 0 | 0 | 0 | | NOTES: ad | 4.3.3.6; 3.5.1.2 | | 0 | 0 | 0 | 0 |
| 13 | antr_dema | anthranila | s_0427[c] | -> | 0 | 0 | 0 | artifical | | | | 0 | 0 | 0 | 0 |
| 14 | N5_antr_s | phosphoril | s_1187[c] | -> | 0 | 0 | 0 | artifical | | | | 0 | 0 | 0 | 0 |
| 15 | carbo_1DF | 1-(2-carbo | s_0076[c] | -> | 0 | 0 | 0 | artifical | | | | 0 | 0 | 0 | 0 |
| 16 | | | | | | | | | | | | | | | |

SRR8994357_WT | SRR8994358_WT | SRR8994359_WT | Common_WT

Fig. 13. Wild type non-flux reaction task result

To perform this test case run the file *TestCase_NonFluxReactions.m* in the 'Scripts' folder of IgemRNA tool.

**b. Filter rate limiting reactions**

Post-optimization task 'filter rate limiting reactions' performs analysis on the generated context-specific models of the same phenotype, the phenotype name is included in the result files (Fig. 14).

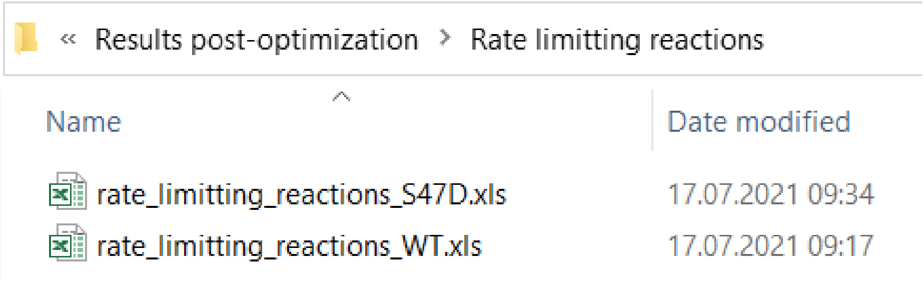


Fig. 14. Rate limiting reactions result folder

Each result file contains sheets for each provided transcriptomics dataset of the same phenotype that has been integrated in the supplied model creating context-specific models. An analysis on these context-specific models have been performed in order to filter reactions that have reached the maximal flux (MaxFlux calculated by FVA) based on the upper bound set according to transcriptomics data and GPR associations. An additional sheet for common rate limiting reactions has also been added to the result file where rate limiting reactions that are present in all datasets are listed (Fig. 15).

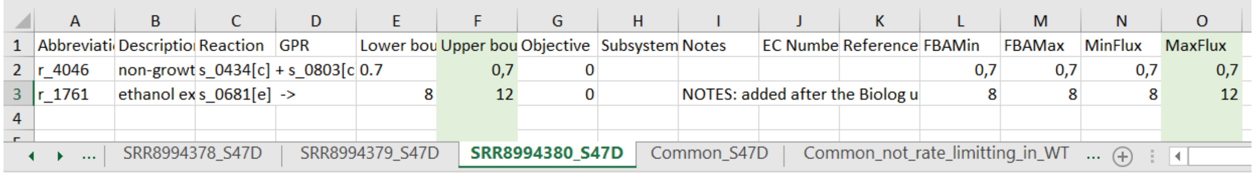

| | A | B | C | D | E | F | G | H | I | J | K | L | M | N | O |
|---|---|---|---|---|---|---|---|---|---|---|---|---|---|---|---|
| 1 | Abbreviati | Descriptio | Reaction | GPR | Lower bou | Upper bou | Objective | Subsystem | Notes | EC Numbe | Reference | FBAMin | FBAMax | MinFlux | MaxFlux |
| 2 | r_4046 | non-growt | s_0434[c] + s_0803[c | | 0.7 | 0,7 | 0 | | | | | 0,7 | 0,7 | 0,7 | 0,7 |
| 3 | r_1761 | ethanol ex | s_0681[e] -> | | 8 | 12 | 0 | | NOTES: added after the Biolog u | | | 8 | 8 | 8 | 12 |
| 4 | | | | | | | | | | | | | | | |

SRR8994378_S47D | SRR8994379_S47D | SRR8994380_S47D | Common_S47D | Common_not_rate_limitting_in_WT ...

Fig. 15. S47D phenotype rate limiting reaction task result

To perform this test case run the file *TestCase_RateLimitingReactions.m* in the ‘Scripts’ folder of IgemRNA tool.

#### c. Flux shifts calculation between different phenotypes

Post-optimization task ‘calculate flux shifts between phenotypes’ compares flux values (calculated by FVA on the context-specific models) between two different phenotypes. In this demonstration flux shifts analysis task was performed on the S47D phenotype datasets using global T1 thresholding approach with the lower global threshold value of 0, these phenotypes were compared to the wild type dataset SRR8994357_WT of the same thresholding approach and threshold values. Result file names include the dataset and phenotype name (provided in transcriptomics data file), used thresholding approach and global threshold values (Fig. 16).

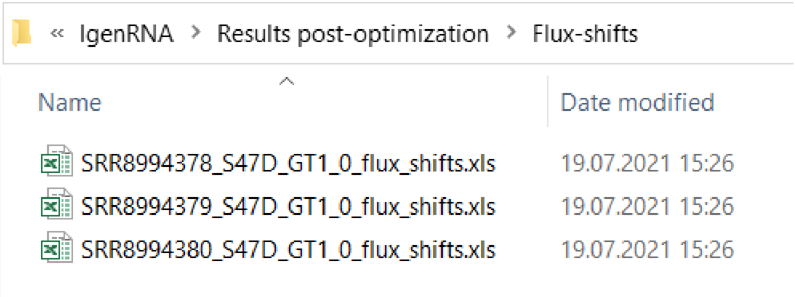


Fig. 16. Flux-shifts result folder

Each result file contains a full reaction list that corresponds to the ‘Reaction List’ sheet in the provided model file as well as additional columns for the calculation results: MinFlux and MaxFlux values (phenotype that is compared, Fig. 17. N, O columns), MinFlux/MaxFlux(dataset name)_(phenotype name)_(thresholding approach)_(global threshold values) phenotype that is used for comparison (Fig. 17. P, Q columns) and the MinFlux/MaxFlux ratio between these two phenotypes (Fig. 17. R, S columns).

| | A | B | C | N | O | P | Q | R | S |
|---|---|---|---|---|---|---|---|---|---|
| 1 | Abbreviation | Description | Reaction | MinFlux | MaxFlux | MinFluxSRR8994357_WT_GT1_0 | MaxFluxSRR8994357_WT_GT1_0 | MinFluxRatio (mutantFlux: wtFlux) | MaxFluxRatio (mutantFlux: wtFlux) |
| 2 | r_0001 | (R)-lactate:fe | s_0025[c] + | 0 | 71.94614 | 0 | 71,94614 | 1 | 1 |
| 3 | r_0002 | (R)-lactate:fe | s_0027[m] + | 0 | 71.94614 | 0 | 71,94614 | 1 | 1 |
| 4 | r_0003 | (R,R)-butane | s_0035[c] + | -11.92121 | 0 | -11,92121 | 0 | 1 | 1 |
| 5 | r_0004 | (S)-lactate:fe | s_0063[c] + | 0 | 63.34479 | 0 | 63,34479 | 1 | 1 |
| 6 | r_0005 | 1,3-beta-gluc | s_1543[c] -> | 0.00748515 | 0.6953993 | 0,007485 | 0,695399 | 1 | 1 |
| 7 | r_0006 | 1,6-beta-gluc | s_1543[c] -> | 0.00250092 | 0.2323451 | 0,002501 | 0,232345 | 1 | 1 |
| 8 | r_0007 | 1-(5-phospho | s_0077[c] -> | 0.000761572 | 8.901205 | 0,000762 | 8,901205 | 1 | 1 |
| 9 | r_0012 | 1-pyrroline-5 | s_0119[m] + | 0 | 74.2016 | 0 | 74,2016 | 1 | 1 |
| 10 | r_0013 | 2,3-diketo-5- | s_0311[c] + | 1.899999e-06 | 16.79361 | 0,000002 | 16,53495 | 1 | 1,015643 |
| 11 | r_0014 | 2,5-diamino- | s_0142[c] + | 1e-05 | 0.0009290385 | 0,00001 | 0,000929 | 1 | 1 |
| 12 | r_0015 | 2,5-diamino- | s_0141[c] + | 1e-05 | 0.0009290385 | 0,00001 | 0,000929 | 1 | 1 |
| 13 | r_0016 | 2-aceto-2-hy | s_0179[m] + | 0.0022135 | 9.177407 | 0,002214 | 9,177407 | 1 | 1 |
| 14 | r_0017 | 2-amino-4-hy | s_0148[m] + | 0 | 5.890104e-05 | 0 | 0,000059 | 1 | 1 |
| 15 | r_0018 | 2-aminoadipa | s_0176[c] + | 0.00328751 | 7.695527 | 0,003288 | 7,695527 | 1 | 1 |
| 16 | r_0019 | 2-dehydropa | s_0149[c] + | 1.899999e-06 | 4.746515 | 0,000002 | 4,746515 | 1 | 1 |
| 17 | r_0020 | 2-deoxy-D-a | s_0552[m] + | 0 | 16.35682 | 0 | 16,35682 | 1 | 1 |
| 18 | r_0021 | 2-hexaprenyl | s_0155[m] + | 0 | 0 | 0 | 0 | 1 | 1 |
| 19 | r_0022 | 2-hexaprenyl | s_0157[m] + | 0 | 0 | 0 | 0 | 1 | 1 |
| 20 | r_0023 | 2-isopropylm | s_0165[c] + | -8.501553 | -0.00340467 | -8,501553 | -0,003405 | 1 | 1 |
| 21 | r_0024 | 2-isopropylm | s_0232[c] + | 0 | 8.442585 | 0 | 8,442585 | 1 | 1 |
| 22 | r_0025 | 2-isopropylm | s_0233[m] + | 0 | 8.501553 | 0 | 8,501553 | 1 | 1 |
| 23 | r_0026 | 2-keto-4-met | s_0294[c] + | 1.899999e-06 | 16.79361 | 0,000002 | 16,53495 | 1 | 1,015643 |
| 24 | r_0027 | 2-methylcitra | s_0834[m] < | 0.00328751 | 7.695527 | 0,003288 | 7,695527 | 1 | 1 |
| 25 | r_0028 | 2-methylcitra | s_0807[m] + | 0 | 25.15836 | 0 | 24,54332 | 1 | 1,025059 |
| 26 | r_0029 | 2-oxo-4-meth | s_0010[c] + | 0.00340467 | 8.501553 | 0,003405 | 8,501553 | 1 | 1 |
| 27 | r_0030 | 2-oxo-4-meth | s_0011[m] + | 0 | 0 | 0 | 0 | 1 | 1 |
| 28 | r_0032 | 3',5'-bisphos | s_0390[c] + | 0 | 69.24784 | 0 | 67,51146 | 1 | 1,02572 |

Fig. 17. Flux-shifts between

To perform this test case run the file *TestCase_fluxShifts.m* in the ‘Scripts’ folder of IgemRNA tool.

Most genome scale metabolic models use biomass objective function to simulate biomass accumulation rates, but in many cases such, *S. cerevisiae* Yeast_8_4 version models have specific wild type function. Optimizing different mutant strains, deletions and specific omics data integration (like transcriptomics data) on the models, can make infeasible optimization solutions although experimental conditions show the opposite. IgemRNA has functionality to remove biomass objective function from model and allow apply flux distribution with transcriptome and/or medium data sets and analyze results.

**d. Nomenclature of file names**

IgemRNA also provides standardized output file naming for easier filtering of analysis datasets (Table 1).

Table. 1

| | **Dataset and Phenotype Name** | **Thresholding Approach** | **Global Threshold Values** |
|---|---|---|---|
| Source | Sheet name in transcriptomics data file (Fig. 1. B) | Based on the selected thresholding approach in the IgemRNA form section B (Fig. 3.) | Based on the provided global threshold values in the IgemRNA form section C (Fig. 3.) |
| Example/Possible values | SRR8994357_WT | A. GT1 (Global T1)<br>B. LT1 (Local T1)<br>C. LT2 (Local T2) | A. 30 (GT1, LT1)<br>B. 30_100 (LT2) |
